# Friction-induced budding of a cancer cell monolayer

**DOI:** 10.1101/2024.02.26.582028

**Authors:** Mathieu Dedenon, Jorge Barbazán, Carlos Pérez-González, Danijela Matic Vignjevic, Pierre Sens

## Abstract

The environment surrounding a tumor plays a crucial role in cancer cell dissemination. Within this microenvironment, cancer-associated fibroblasts (CAFs) generate compressive forces and actively remodel tumors. Using *in vitro* circular clusters of cancer cell monolayers surrounded by CAFs, we generate structures that are reminiscent of multicellular buds observed in vivo for colo-rectal cancer. A supracellular contractile ring spontaneously assembles at the inner edge of the CAF monolayer and drives its closure on top of the cancer cells through a purse-string mechanism. The frictional shear stress exerted by CAFs triggers multilayering of cancer cells, followed by the emergence of a multicellular bud constricted by the CAF ring. To explain this observation, we developed a theoretical model based on continuum mechanics. This model outlines the early transformations in the shape of cancer cell monolayer and links the layering of cells to a general criterion involving height deformation. It identifies the specific physical conditions that favors budding, and reproduces the observed dependence of the budding probability and bud sizes with the diameter of the cancer cell cluster. Our findings highlight the importance of active mechanical interactions between the tumor and its micro-environment on aggressive modes of cancer invasion.

## Introduction

Tumor invasion is a critical stage of tumor development that initiates the metastasis process, significantly influencing patient survival. Cancer cells disseminate from the primary tumor either as single cells, through a process called epithelial to mesenchymal transition(1–3), or by the detachment of small cell clusters through the process called tumor budding(4). These multicellular buds are notably linked to a higher metastatic occurrence and a decrease in patient survival rates(5, 6). Cancer invasion is not a cell-autonomous process; the tumor micro-environment, including various types of stromal cells and the extracellular matrix, plays an important role in cancer dissemination. Cancer-associated fibroblasts (CAFs) are key players in cancer progression(7, 8). They promote tumor dissemination by breaking the basement membrane (9), remodeling extracellular matrix (10, 11), leading migration of cancer cells(12, 13), and secreting growth factors that modulate cancer cell behaviour and the infiltration of immune cells(14). However, due to their highly contractile nature, CAFs can also act in a protective manner by encapsulating the tumor, potentially hindering its spread(15–17).

Tumor budding is especially common in colorectal cancer(4), a type of cancer where CAFs are particularly abundant(18). In the present work, we investigate the role of CAFs on “tumor budding” in an idealised *in-vitro* context and propose a theoretical model based on frictional forces between tissues in relative motion.

### Experimental system

To mimic the tumor and its micro-environment in a controllable manner, we designed a 2D *in vitro* system made of primary Cancer Cells (CCs) surrounded by CAFs (Fig. 1A). The experimental setup and its precise characterisation are described in detail in (17). The CCs are initially patterned to form a cluster of a controlled radius. CAFs are later seeded to surround the cancer cells (Fig. 1B). After a phase of proliferation, CAFs surrounding the CC cluster align tangentially to its periphery and spontaneously assemble a supracellular contractile ring enriched in actin and myosin II (Fig. 1A, Left and (17)). Ring contraction triggers the closing of the CAF monolayer on top of the CC cluster (Fig. 1A, Middle) through a purse string mechanism also seen in epithelial gap closure experiments (19–21). Ring closure may be successful, with the CAF monolayer completely covering the CC cluster, or incomplete, when the ring stalls around a bulge of cancer cells (Fig. 1B,C and movie S1). The latter originates from a 3D reorganization of CCs into a multi-cellular ‘bud’ which resists ring compression (Fig. 1A, Right and (17)). This phenotype can be seen by quantifying the evolution of the ring radius *R*_*c*_(*t*) with time for each CC cluster (Fig. 1D). After an initial phase of closure with a constant velocity of order *V*_*c*_ ≡ |d*R*_*c*_*/*d*t*| ∼ 3*µ*m/hr, *V*_*c*_ decreases with a quasi-stabilization of the ring radius *R*_*c*_(*t*) after one or two days (Fig. 1D).

**Fig. 1.**
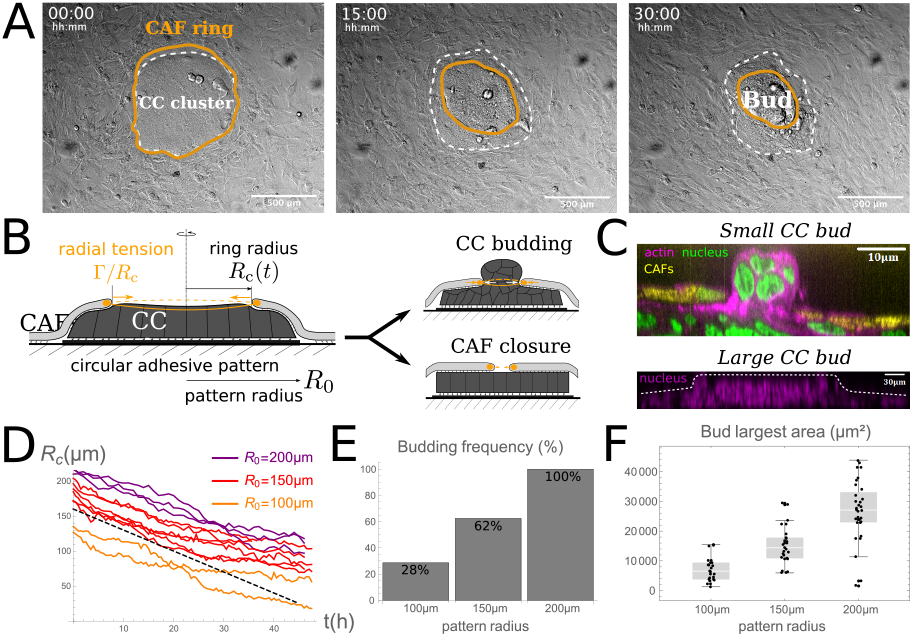
Experimental system. **(A)**: Phase contrast images of the experimental co-culture, made of a monolayer of Cancer Cells (CCs) with controlled size surrounded by Cancer-Associated Fibroblasts (CAFs), at three different times from left to right. CAFs assemble a supracellular contractile ring at the edge of CCs (left), climb on top of CCs to undergo gap closure (middle), and undergo either full closure or stall around a CC bud (right). *R*_0_ = 150*µ*m. See supplementary movie S1. **(B)**: Sketch of a side view of the CAF-CC system (Left) showing CAFs on top of CCs, undergoing gap closure thanks to the ring (orange) with instantaneous radius *R*_*c*_(*t*). The system can evolve into CC budding phenotype at steady-state (Top right), or into full CAF closure (Bottom right). **(C)**: Microscopic fluorescent image of the system where a small (top) or large (bottom) bud has emerged from 3D cell rearrangements and arrests CAF closure, for *R*_0_ = 150*µ*m. **(D)**: Evolution of the CAF ring radius *R*_*c*_(*t*) for different clusters with CC budding phenotype (n=11). The dashed line indicates a constant CAF velocity *V*_*c*_ = |d*R*_*c*_*/*d*t*| = 3*µ*m/h, a typical initial magnitude for all clusters. **(E)**: CC budding frequency as a function of pattern radius, observed for *R*_0_ = 100 (n=14), 150 (n=8) or 200 (n=7) microns. **(F)**: Bud projected area as a function of pattern radius, observed for *R*_0_ = 100*µ*m (n=24), 150*µ*m (n=27) or 200*µ*m (n=32). The area is measured from a top view at steady-state.

A natural control parameter to understand the appearance of the budding phenotype is the size of the cancer cell cluster (Fig. 1B). Indeed among a population of clusters, the experimental budding frequency is found to be correlated to the pattern size (Fig. 1E). In addition at steady-state, the projected area of buds also appears correlated with pattern size despite some large dispersion on individual clusters (Fig. 1F). These two features will serve as important experimental tests for the theoretical model of the early stage of CC budding proposed in this paper, which culminates into phase diagrams describing the budding likelihood.

### Theoretical model

The driving force for the budding process is the CAF contractile ring, modelled here as a 1D circular cable of radius *R*_*c*_ and a line tension Γ. This generates a compressive radial tension Γ*/R*_*c*_ by Laplace law ((17) and Fig. 1B). The ring line tension (Γ ∼ 0.5 − 1*µ*N) has been measured by replacing the CC clusters with elastic pillars of known stiffness (17). Although the actomyosin ring does not apply direct pressure on the CCs prior to multilayering, the relative motion of the CAF layer on top of the CC cluster generates a frictional shear which drives the compression of the CC cluster. The shear stress is proportional to the velocity difference: *τ* = *ξ*(*v*_CAF_ −*v*_CC_) with a friction coefficient *ξ* (Fig. 2A, Left). If no such mechanical interaction exists (*ξ* = 0), the two tissues evolve independently with the CAFs closing on top of an unperturbed CC cluster. Friction creates inward CC compression which may lead to budding. This morphological transition requires 3D cell rearrangements, where some cells loose their basal connection with the substrate. Because the experimental population of clusters nicely separates in two phenotypes (Fig. 1B,D,E), it seems reasonable to assume a strict causal link between the triggering of CC multilayering and the budding phenotype. Starting from a weakly deformed CC monolayer (Fig. 2A, Left), multilayering events are assumed to occur when cells reach a critical height *h*^*^ (Fig. 2A, Middle), which then triggers a cascade of events from a localized proto-bud (Fig. 2A, Right) to the late budding state. We assume rotational symmetry (a circular ring) for simplicity and focus on the early evolution of the monolayer height field *h*(*r, t*) (Fig. 2B). Calling respectively *R*_*T*_ (*t*) and *R*_*c*_(*t*) the instantaneous radii of the CC monolayer and CAF ring, the system is separated in three spatial domains (Fig. 2B): the CAF-free region (0 *< r < R*_*c*_), the CAF-covered region (*R*_*c*_ *< r < R*_*T*_) and the surroundings (*r > R*_*T*_). Here, we neglect cell divisions of CCs and assume cell incompressibility, so that monolayer deformation leads to a displacement of peripheral radius *R*_*T*_ (*t*).

**Fig. 2.**
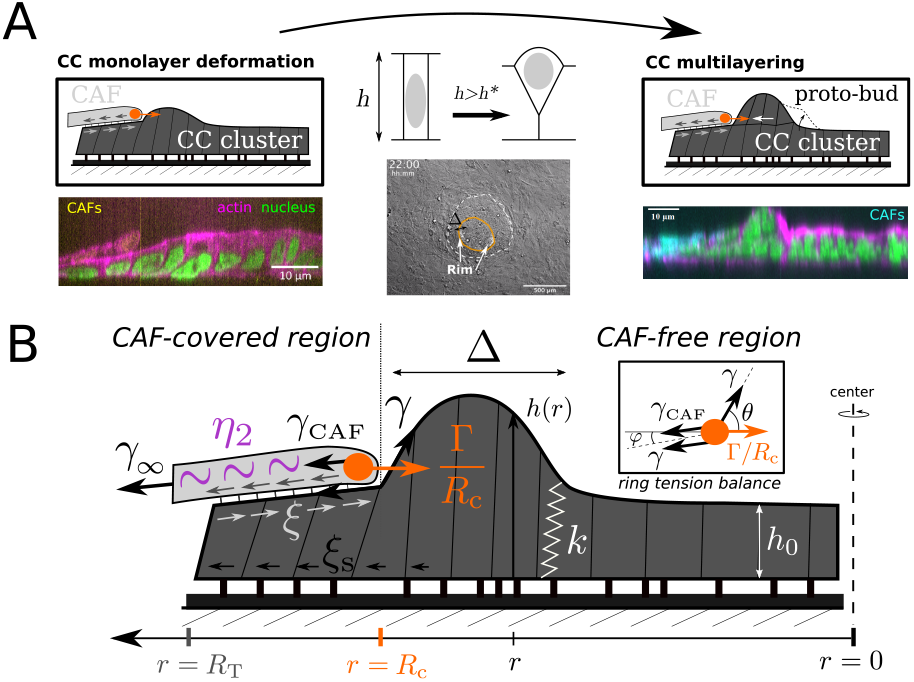
Theoretical model. **(A)**: Left: Sketch and microscopic fluorescent image of the CAF-CC system when the CC cluster is weakly deformed as a monolayer. Middle: The loss of basal interface is assumed to be triggered when the cell height *h* is above the critical value *h*^*^ (Top). The monolayer deformation tends to be local-ized spatially near the CAF ring, into a rim of width Δ (Bottom and **movie**). Right:Sketch and microscopic fluorescent image after few CC multilayering events, showing the localised rim of deformation). **(B)**: Sketch for the theoretical model of bud initiation showing a deformed CC monolayer with height field *h*(*r*) and peripheralradius *R*_*T*_, the CAF layer with tension field *γ*_CAF_(*r*) and surrounding pre-tension*γ*_∞_ = *γ*_CAF_(*r* = ∞), the actomyosin ring at radius *R*_*c*_ with line tension Γ. CCmechanics is described through an elastic stiffness *k* and equilibrium height *h*_0_ and an apical tension *γ*. The tensions are mechanically equilibrated at the CAF ring (inset). Dynamic friction occurs at CC-substrate and CAF-CC interfaces, withrespective coefficients *ξ*_s_ and *ξ*. This ensures the localization of CC deformation in a rim of width Δ on the CAF-free region.

We adopt the framework of continuous mechanics to derive equations for the evolution of the local CC height (see S.I. for details). We assume an elastic behaviour for the CC monolayer. Deviation from a natural height *h*_0_ causes an intracellular pressure associated to a stiffness *k*. The total pressure also includes the Laplace pressure coming from the curvature *C*_free_ of the apical surface of the CCs. On the CAF-free region, the apical tension is written *γ* and the constitutive equation for the pressure field reads:

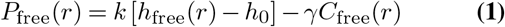

with the apical curvature expressed in the small deformation limit as: 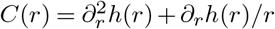. This equation is shown to be consistent with the mechanics of a bubbly vertex model in the weak deformation limit, as detailed in the supplementary Information [S.I.].

In the CAF-free region, the flow of CCs over the substrate is associated with pressure gradients. We consider a generic frictional stress between the CCs and the substrate proportional to CC velocity field *υ*_free_(*r*), with a friction coeffi-cient *ξ*_s_ (Fig. 2B). To least order in the CC deformation (|∂_*r*_*h*| *≪* 1), the horizontal force balance reads (see S.I. for details)

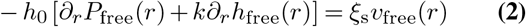

and local volume conservation imposes ∂_*t*_*h*_free_ = −*h*_0_∂_*r*_ [*rυ*_free_] */r*. Eliminating pressure and velocity fields yields a PDE for the height field *h*_free_(*r, t*) on the CAF-free region

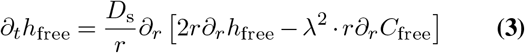

with a diffusion constant *D*_s_ ≡ *k*(*h*_0_)^2^*/ξ*_s_ and a mechanical length 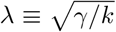. The shape of the CAF-free CC region thus obeys a diffusion-like equation with a moving boundary (the CAF ring), which leads to the formation of a localised rim of width Δ near the moving boundary, consistent with our observations (Fig. 2A). This localization is reminiscent of the rim formation occurring upon liquid film dewetting (22– 24), although in that case, the rim width is controlled by the slip length *b* ≡ *η/ξ*_s_ associated with film viscosity *η* and film-substrate friction *ξ*_s_.

On the CAF-covered region, the thin CAF layer is described as a continuous surface that follows the CC monolayer shape. Its tension *γ*_CAF_(*r*) varies in space because of CAF-CC friction (Fig. 2B), and local force balance gives ∂_*r*_*γ*_CAF_(*r*) ≃ *ξ* [*υ*_CAF_(*r*) − υ_cov_(*r*)] for weak CC deformations. Assuming surface incompressibility for CAFs, one can directly obtain the CAF lowest order velocity field *v*_CAF_(*r, t*) = −*V*_*c*_(*t*)*R*_*c*_(*t*)*/r* from the closure velocity *V*_*c*_(*>* 0). The rate of gap closure results from a balance between the driving force (the ring line tension Γ) and dissipation. Possible mechanisms are discussed in the S.I.. We assume here that dominant dissipation comes from the CAF tissue viscosity *η*_2_ and that dissipation from CAF-CC friction *ξ* can be treated as a small perturbation. This choice is motivated by the experimental closure dynamics (Fig. 1D), where *V*_*c*_(*t*) appears approximately constant - independent of the initial CC size - at early times. Indeed for *ξ* = 0, one can show that *V*_*c*_(*t*) = Γ*/*(2*η*_2_). Knowing orders of magnitude for both velocity and line tension, one gets *η*_2_*/h*_CAF_ ∼ 107 − 109Pa.s when dividing by CAF thickness *h*_CAF_, a reasonable value for this type of tissues (25–28).

On the CAF-covered region, normal force balance includes the CAF tension, and tension anisotropy due to CAF viscosity: *γ*_*rr*_(*r*) = *γ* + *γ*_CAF_(*r*) + 2*η*_2_*R*_*c*_*V*_*c*_*/r*^2^ and *γ*_*θθ*_(*r*) = *γ* + *γ*_CAF_(*r*) − 2*η*_2_*R*_*c*_*V*_*c*_*/r*^2^. Eq. 1 thus becomes:

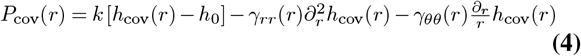

The friction term modifies the horizontal force balance as:

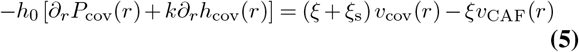

Eliminating the velocity and pressure fields to obtain a single PDE for the height field can be done as above in the small friction limit to obtain a final growth PDE for *h*_cov_ to least order in *ξ* (see S.I. Eq.S17).

Force balance at the boundary between CAF-covered and CAF-free regions (*r* = *R*_*c*_) imposes a discontinuity of the slope ∂_*r*_*h*. Vertical force balance reads *γ*_*rr*_ sin(*ϕ*) = *γ* sin(*θ*) (angles are defined in Fig. 2B). Horizontal force balance involves the driving force for CAF closure Γ*/R*_*c*_ and controls ring dynamics: for a flat CC cluster mechanically decoupled from CAFs (*ξ* = 0), it reads *R*_*c*_*γ*_CAF_(*R*_*c*_) + 2*η*_2_*V*_*c*_ = Γ. Finally on the surrounding region, we describe the CAF tissue as a simple sheet under active tension *γ*_∞_ (Fig. 2B). Amore complex CAF description was proposed in (17) to account for the traction force pattern on the substrate, but the present model is sufficient here as we focus on the CC cluster mechanics. The active tension modifies the (friction-free) closure dynamics according to *V*_*c*_ = (Γ − *γ*_∞_*R*_*c*_)*/*(2*η*_2_). As the ring driving force decreases with the cluster size, there exists a maximal cluster initial radius *R*_0_ = Γ*/γ*_∞_ beyond which the ring does not close. With Γ ∼ 0.5 − 1*µ*N (17) and *γ*_∞_ ≲ − 3mN/m (29), CAF closure is expected for *R*_0_ ;S 500 − 1000*µ*m. Indeed we observe CAF closure initiation for all clusters of the largest pattern radius (*R*_0_ = 200*µ*m), which implies *γ*_∞_ *<* 5mN/m.

## Results

The theoretical description of the CAF-CC system consists in the resolution of two growth PDEs for the CC monolayer height field *h*(*r, t*) on evolving domains [0; *R*_*c*_(*t*)] and [*R*_*c*_(*t*); *R*_*T*_ (*t*)], with initial conditions *h*(*r, t* = 0) = *h*_0_ and *R*_*c*_(0) = *R*_*T*_ (0) = *R*_0_. Spatial boundary conditions are detailed in the S.I.. Parameters are of mechanical (*k, γ, γ*_∞_, *Γ*), dissipative (*η*_2_, *ξ, ξ*_s_) and geometric (*h*_0_, *h*^*^, *R*_0_) nature. The typical CC monolayer height is of order *h*_0_ *∼* 10*µ*m (see S.I.). CAF-CC friction can be treated as a perturbation for closure dynamics provided 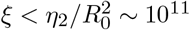 which is expected for such systems (19, 27, 28, 30). We eliminate *h*_0_, Γ and *η*_2_ to make all quantities dimensionless and use the same notation for the remaining parameters.

### Flat covered limit

To assess the main features of the model, we first consider a simplified case where the CAF-covered region remains flat (Fig. 3): *h*_cov_(*r, t*) = *h*_0_. This corresponds to the formal limit where Γ, *γ*_∞_, *η*_2_ → ∞, while the initial closing velocity *V*_*c*_ = (Γ − *γ*_∞_*R*_*c*_)*/*(2*η*_2_) remains finite (see S.I.). The CAF frictional shear stress drives an inward displacement of the CCs (*V*_*T*_ *>* 0) which acts as a source of CC deformation in the CAF-free region (Fig. 3A). In the CAF-covered region, both velocity fields are determined by the boundary dynamics: *v*_cov_(*r*) = −*V*_*T*_ *R*_*T*_ */r* and *v*_CAF_(*r*) = −*V*_*c*_*R*_*c*_*/r*. The integration of Eq. 5 gives a pressure field *h*_0_*P*_cov_(*r*) = [*ξR*_*c*_*V*_*c*_ − (*ξ* + *ξ*_s_)*R*_*T*_ *V*_*T*_] log(*R*_*T*_ */r*) under the CAF layer.

**Fig. 3.**
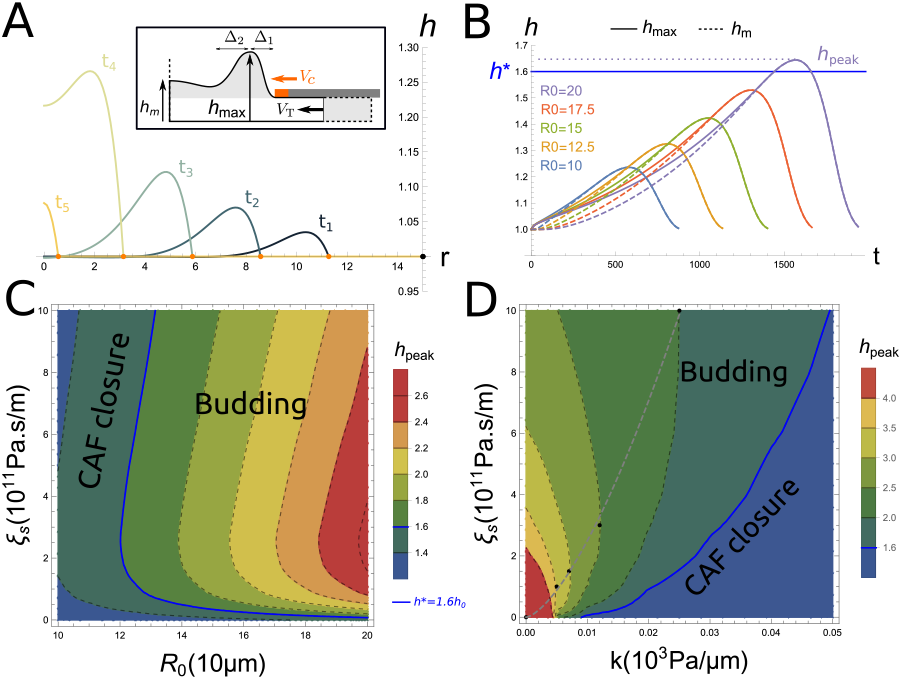
Results of the flat covered limit (no deformation on the CAF-covered region): **(A)**: Snapshots of the CC monolayer shape *h*(*r, t*) at different times for *ξ*_s_ = 2, *R*_0_ = 15. The CC peripheral radius *r* = *R*_*T*_ is the black dot and the ring position *r* = *R*_*c*_ is the orange dot (see supplementary movie S2). *Inset* : Sketch of the system. **(B)**: Evolution of the maximal height *h*_max_(*t*) and the central height *h*_*m*_(*t*) with time for different pattern radii *R*_0_, for *ξ*_s_ = 0.1. At late time, the rim disappears (*h*_max_ = *h*_0_) and a maximum deformation *h*_*peak*_ is reached, correlated with pattern radius *R*_0_. **(C)**: Phase diagram of *h*_*peak*_ in the parameter space (*R*_0_, *ξ*_s_). A critical height *h*^*^ = 1.3 defines the CC budding threshold (blue line). **(D)**: Phase diagram of *h*_*peak*_ in the parameter space (*k, ξ*_s_), for *R*_0_ = 20. Black points indicate the maximal deformation at 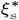. The gray dashed line shows the predicted maximal deformation assuming it occurs when elastic and Laplace pressures are comparable. The height *h* and *r* are in unit *h*_0_ = 10*µ*m. Other parameters are *k* = *γ* = 0.01, *γ*_∞_ = 0, *ξ* = 0.1.

In the flat covered limit, the displaced peripheral CC volume is entirely transferred into the deformation of the CAF-free region at a rate d_*t*_ν_up_ = 2*πh*_0_*R*_*T*_ *V*_*T*_. The CC deformation is localised near the CAF ring due to friction with the substrate, forming a rim of width Δ(*t*) that initially grows as Δ(*t*) ∼ (*λ*^2^*D*_s_*t*)1*/*4 (Eq. 3) before reaching a plateau value Δ_*c*_. Two length scales then characterise the deformed shape: a mechanical length 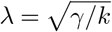 and a dynamic length *λ*_*V*_ = *D*_*s*_*/V*_*c*_ = *kh*_0_^2^*/*(*ξ*_*s*_*V*_*c*_). The rim is asymmetric ((Fig. 3A)) and a scaling analysis presented in the S.I. shows that it is characterised by two length scales: Δ_*c*,1_ ∼ (*λ*^2^*λ*_*V*_)^1/3^ and Δ_*c*,2_ ∼ (*λλ*_*V*_)^1/2^ for finite *λ*. In addition, one gets Δ_*c*_ ∼ *λ*_*V*_ in the limit *γ* → 0 and Δ_*c*_ ∼ (*λ*^2^*λ*_*V*_)^1/3^ in the limit *k* → 0. Increasing the CC substrate friction *ξ*_*s*_ results in a more localised deformation, i.e. a smaller rim width. At longer times, when *R*_*c*_(*t*) ≃ Δ_*c*_, the CC rim merges into a bump covering the entire CAF-free region (Fig. 3A).

Fig. 3B shows the evolution of the maximum CC deformation (the top of the rim) with time *h*_max_(*t*). It exhibits an initial rise up to a peak value *h*_*peak*_ (Fig. 3B), followed by a relaxation to its initial value *h*_0_. The central height of the CC cluster *h*_*m*_(*t*) is also shown. It remains unperturbed as long as the rim is spatially localized, and grows towards *h*_max_ when the annular rim merges into a bump for *R*_*c*_(*t*) ≃ Δ_*c*_ (Fig. 3A,B). The existence of a peak deformation indicates that the initial CC flow caused by CAF-CC friction reverses to an outward back-flow when the CC pressure overcomes the frictional driving force. While the end state predicted by the model is always a closed CAF layer on top of an undeformed CC cluster, we assume that irreversible multilayering and the initiation of the budding phenotype occurs when the CC peak height *h*_*peak*_ reaches a threshold distributed around a critical value *h*_*crit*_. Therefore, the system’s final state can be understood from the study of how *h*_*peak*_ depends on the different parameters.

First, Fig. 3B shows that the peak height increases with the pattern radius *R*_0_. This behaviour is recovered in the limit of small substrate friction: *D*_*s*_ ≫ *V*_*c*_*R*_0_, where the shape of the CAF-free region can be calculated analytically. It predicts *δh*_*peak*_ ∝ *ξ* Γ*R*_0_*/*(*kη*_2_) for small tension *λ ≪ R*_0_, or 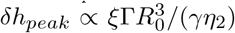 for large tension *λ ≫ R*_0_ (see S.I.). As expected, the deformation increases linearly with the CAF-CC friction coefficient *ξ* and decreases with the CC mechanical resistance (*k* and *γ*). The correlation between system’s size and CC deformation agrees with the experimental observation and will be studied in more details below. A rather non-trivial result is the influence of substrate friction on CC deformation, recapitulated in two phase diagrams (Fig. 3C,D).

The first phase diagram, in the (*R*_0_, *ξ*_s_) parameter space (Fig. 3C), confirms that the peak deformation correlates with *R*_0_, but also shows that the CC-substrate friction parameter *ξ*_s_ has a non-monotonic influence on the deformation peak. A maximal CC deformation occurs for a finite value of 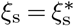. This non-monotonic feature is the result of several antagonistic effects. First, substrate friction limits the inward flow of CC driven by CAF friction (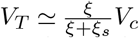 at early times), which limits the volume transferred from the covered to the free region. Second, the CC back-flow at later stage is hindered by substrate friction, which tends to increase the peak deformation. Finally, the deformation is localised by substrate friction (the rim width Δ decreases with increasing *ξ*_*s*_), which concentrates deformation, but also increases the rim pressure driving the back-flow.

The relative importance of the different effects is studied in the S.I. using toy models of simplified geometries. They show that the peak deformation monotonically increases with substrate friction in the absence of localisation or if the Laplace pressure is neglected (*γ* → 0), but monotonically decreases with substrate friction if the back-flow is driven solely by Laplace pressure (*k* → 0). These results are confirmed by the second phase diagram in the (*k, ξ*_s_) parameter space (Fig. 3D), which shows a monotonic decrease of the deformation with substrate friction for small *k* and a monotonic increase for large *k*. The largest deformation is obtained for a particular value of substrate friction 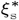 which increases with the stiffness *k*. This optimum corresponds to the parameters for which the elastic pressure *kδh* and the Laplace pressure ∼ *γδh*″ play comparable role in driving the back-flow, that is *λ* ∼ *λ*_*V*_ or 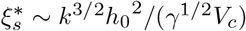 (gray dashed line in Fig. 3D).

### Deformable CAF layer

In S.I., we show using a toy model that the “flat covered” limit is approximately valid when Γ *≫ γR*_0_(1 + (*R*_0_*/λ*)^2^). In both cases, this requires small surface tension *γ ≪* 10^−2^N/m and small CC stiffness *E* ≡ *k*.*h*_0_ *≪* 10Pa. The second condition is unlikely to be valid for the *in vitro* CAF-CC system, since one expects CC stiffness values *E* ∼ 0.1 − 10kPa (31). It is thus necessary to consider the deformation of the CAF-covered region, obtained by enforcing local force balance throughout the CC cluster (Eq. 5). Numerical results accounting for CAF deformation are shown in Fig. 4. CAF deformation is a significant fraction of the maximum deformation when CAF pre-tension is small (*γ*_∞_ → 0, Fig. 4A). If the substrate friction *ξ*_s_ is sufficiently large (Δ_*c*_ *≪ R*_0_), a rim develops ahead of the closing CAF ring as discussed before. The deformation then evolves in two qualitatively distinct fashions depending on the parameters (see sketches in Fig. 4B). For small substrate friction, the rim persists until it merges into a bulge and the situation is qualitatively similar to the “flat covered” case. For large substrate friction, the rim becomes covered by the CAFs. This is illustrated by the evolution of the maximum height (*h*_*max*_) and the ring height (*h*_*c*_) with time (Fig. 4B). The former is consistently higher than the latter for small substrate friction (left panel) indicating the persistence of the free rim, while the two heights are similar except near ring final closure for high substrate friction, indicating that the rim is covered by CAFs (right panel). More examples of temporal evolution in the two regimes are shown in the S.I..

**Fig. 4.**
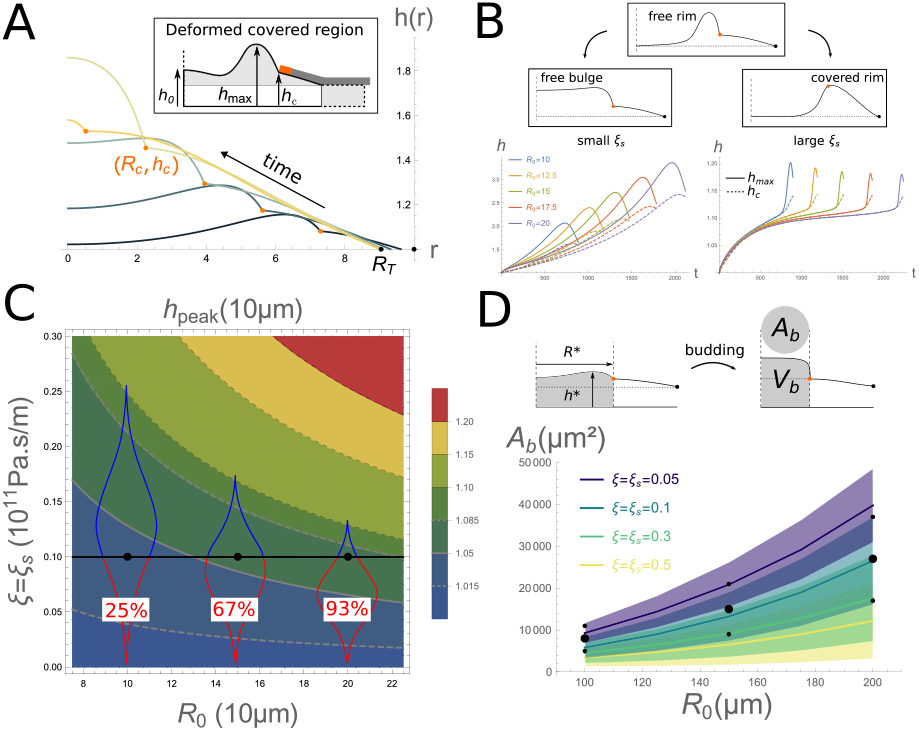
Results with deformable covered region: *k* = 0.01 *(A-B), k* = 0.05 *(C-D), γ* = 0.01, *γ*_*∞*_ = 0, *ξ* = 0.5. **(A)**: Snapshot of the temporal evolution of the CC monolayer shape with height *h*(*r, t*), CC peripheral radius *R*_*T*_ (black point) and ring position (*r* = *R*_*c*_, *z* = *h*_*c*_) (orange point). *ξ*_s_ = 0.5, *R*_0_ = 10 (see supplementary movie S3). **(B)**: A free rim can evolve into a free bulge for low substrate friction (Left, *ξ*_s_ = 0.1) or a covered rim for high substrate friction (Right, *ξ*_s_ = 5). **(C)**: Phase diagram of the maximal height *h*_peak_ in parameter space (*R*_0_, *ξ*) at *ξ*_s_ = *ξ. h*^*\**^ has a normal distribution with mean indicated by gray line and two s.d. by dashed gray lines. Budding occurs when *h*_max_(*t*) *< h*^*\**^, and probabilities are indicated for *R*_0_ = 10, 15, 20. **(D)**: Theoretical and experimental bud projected area *A*_*b*_ for the three available pattern radii *R*_0_. Like in (C), one uses a critical strain *ϵ* ^*\**^ *≡* (*h*^*\**^ *− h*_0_) *≃ h*_0_ [0.7, 5, 16, 27]% for *ξ* = *ξ*_s_ = 0.05 : 0.5. One assumes the final bud to be cylindrical with two cell layers such that its height is *h*_*b*_ = 2*h*_0_ (top). Using *V*_*b*_ = *πh*_0_(*R*^*\**^)^2^ = *A*_*b*_.*h*_*b*_, one can compute a theoretical projected area with *R*^*\**^ = *R*_*c*_[*h*_max_ = *h*^*\**^].

Unlike in the flat covered regime, the deformation does not vanish at the moment of ring closure, since it can be accommodated by the deformability of the CAF layer. There is nevertheless a clear peak deformation at a given time, confirming the existence of a CC back-flow. The system continues to evolve after CAF closure and eventually returns to the undeformed state.

In the experiments, we observe a significant inward flux of CC, *V*_*T*_ *≃ V*_*c*_*/*2, at the onset of ring closure (see S.I. Fig.S2), which suggests that the two friction coefficients are of the same order (*ξ ≃ ξ*_*s*_). Fig. 4C shows that in this regime, the peak height *h*_*peak*_ increases both with the cluster size *R*_0_ and the friction parameters. Interestingly, the correlation between cluster size and CC deformation is not necessarily observed for large substrate friction *ξ*_*s*_ *≫ ξ*, in which case most of the CC deformation is covered by CAFs (the covered rim regime - left panel of Fig. 4B) and the maximum CC deformation *decreases* with increasing cluster size. This feature is further discussed in the S.I. but is not believed to be relevant to our experimental situation.

The correlation between cluster size and peak CC deformation for small surface friction suggests that larger clusters should be more likely to exhibit CAF-driven budding, as observed experimentally (Fig. 1E). To quantitatively assess the budding probability, we postulate that budding is triggered when the cell strain *ϵ* = (*h*_*max*_ *−h*_0_)*/h*_0_ reaches a threshold value *ϵ* ^*\**^, assumed to be gaussian distributed about a mean value due to cell variability. Since the peak deformation increases with friction, the distribution of strain thresholds corresponds to a distribution of friction thresholds for each cluster size, represented as violin plots on the contour plot Fig. 4C. This allows to calculate the budding probability for a given size as the fraction of the population whose threshold friction is below the actual friction value. The model should also be able to reproduce the observed distribution of bud sizes as a function of the cluster size (Fig. 1F). To obtain this, we postulate that if the budding threshold is met at a given time *t*^*\**^: *h*_max_(*t*^*\**^) = *h*_0_(1 + *ϵ* ^*\**^), the volume of the resulting bud is the volume of the free CC layer at the budding threshold: *V*_*b*_ = *πh*_0_*R*_*c*_(*t*^*\**^)^2^. A knowledge of the bud shape is required to convert the volume into the experimentally accessible bud projected area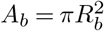. While small buds can be equally described with a spherical or cylindrical geometry, both experimental observation and theoretical analysis on large buds favour a roughly cylindrical shape (Fig. 1C) with a height *h*_*b*_ estimated from confocal microscopy to be a few cell height: *h*_*b*_ *∼* 2 *−* 4*h*_0_ ((17) and S.I.). Assuming *h*_*b*_ to be independent of the bud radius *R*_*b*_ for simplicity leads to a bud area *A*_*b*_ = *V*_*b*_*/h*_*b*_ that can be compared to experiments, as show in Fig. 4G.

Obtaining an appropriate fit for both the budding probability and the resulting bud size strongly constrain the model parameters. The experimental budding probability (Fig. 1F) shows an almost linear dependence on the cluster size. This precludes large friction values, for which the budding probability shows a stronger dependence on cluster size (Fig. 4C). Furthermore, large friction coefficients induce in more localised deformations. Consequently, the peak deformation which is close to the threshold *h*_peak_ *∼ h*^*\**^ for intermediate cluster sizes, as the budding probability is closed to one half occurs for a smaller ring radius. This means that larger friction coefficients result in small bud sizes incompatible with our observations (Fig. 4G). Small buds are also obtained if the CC mechanical resistance is weak (small *k* and/or *γ*), because the back flow responsible for the deformation peak occurs for small ring radii (see S.I.). As shown in Fig. 4C, the relevant regime corresponds to the small friction case with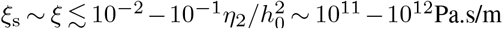, for which the perturbative approach is marginally valid.

## Discussion

Using an *in vitro* co-culture of a patterned cluster of cancer cells (CC) surrounded by CAFs, we show that mechanical interactions between the two cell types is sufficient to generate rearrangement of the cancer cells leading to long-lived bud-like structures. This CC budding phenotype originates indirectly from the contractility of a supracellular actomyosin ring that spontaneously assembles at the inner periphery of the CAF monolayer, and drives its closure on top of the CC cluster in a purse string mechanism.

We develop a theoretical model of this process based on the transmission of the mechanical stress generated by CAF closure to the cancer cells via a generic friction, which could originate from non-specific as well as specific interactions. The latter includes heterotypic binding of E-cadherins and N-cadherins (13), or the matrix secreted by the CAFs (32). Our novel theoretical approach adopts a continuous elastic description of the CC cluster as an epithelial tissue on a substrate that combines in-plane and out-of-plane elastic stresses, and includes viscous dissipation. Comparison with a bubbly side vertex model of a cell monolayer shows that our continuous approach appropriately describes the balance of forces at the lateral interfaces of cancer cells, and the modulation of cell height resulting from intracellular pressure gradients. The latter is central to the initiation of 3D cell rearrangements such as those accompanying monolayer budding. Here we restrict ourselves to friction-dominated CC dissipation. Cell rearrangements within the monolayer could be included as viscous dissipation in our continuous equations (33).

The model shows that a generic dynamic friction between the CC and the CAFs is sufficient to explain important experimental features of the budding phenotype. The friction-driven CC inward flow accompanying CAF closure leads to a build-up of intracellular CC pressure and an increase of cell height. The pressure is maximum close to the CAF contractile ring, leading to the formation of a deformed CC rim as observed experimentally. The budding probability and the geometric features of the cancer buds may then be obtained by invoking the existence of a stress or strain threshold for CC multilayering.

Despite its simplicity, our conceptual approach leads to a surprisingly complex theoretical state space. Specifically, the observed correlation between CC budding frequency and pattern size is non trivial. The CAF-CC and substrate-CC frictions parameters are critical for this behaviour. The former is responsible for the transmission of stress between the two cell types, and the latter strongly affects the spatial distribution of intracellular CC pressure and controls both the appearance and the persistence of the CC rim. This behaviour is also modulated by CC stiffness and CC surface tension. Experimental observations correlating bud properties to pattern size highly constrain the parameter space to specific regions of relevance. For instance, if the CC-substrate friction is too weak, no rim forms and the CC deformation spreads over the entire CAF-free region. Budding probabilities compatible with experimental data then require unreasonably small values of the critical strain for multilayering. If the CC-substrate friction is too large, the model predicts bud sizes that are too small, and possibly a negative correlation between the maximum CC deformation and the cluster size, inconsistent with the observed budding probabilities. Similarly, weak CC stiffness or surface tension predict too small buds whereas too large values prevent monolayer deformation.

One strength of our *in-vitro* approach is the ability to control geometric parameters such as the CC cluster shape and size. This is afforded by the presence of a substrate, which challenges the direct applicability of our model to physiological *in vivo* tumors interacting with their environment. CAFs have been shown to compress cancer cells *in vivo* using actomyosin contractility, leading to tumor compartmentalization (17). However, a real tumor would be better described by a CC spheroid surrounded by extracellular matrix and CAFs (10). Whether the CAF organisation could be appropriately described as a monolayer in this context is unclear, but the friction-induced shear stress between CAFs and cancer cells would nevertheless remain an important component of the mechanical interaction between the two cell types, in addition to other mechanisms such as direct infiltration within the tumor. On the other hand, the friction with the substrate, which we identify as an important parameter controlling budding *in vitro*, is absent in spheroids and should be replaced by viscous-like dissipation associated with cell rearrangements within the tumor. Therefore, a direct extension of this work will be to focus on the experimental and theoretical characterization of this 3D *in vitro* system.

## ACKNOWLEDGEMENTS

We acknowledge funding from: INSERM ITMO cancer 20CR110-00 (DV, PS), H2020 Marie Skłodowska-Curie grant agreement 659776 (JB), Fondation ARC pour la Recherche sur le Cancer (CPG), ANR-11-LABX-0038 (MD, JB, CPG, DV, PS)

## SI Movies

Movie S1: Real time evolution of *in-vitro* cancer cells-CAFs co-cultures (Fig. 1A). Movie S2: Theoretically computed temporal evolution of the CC shape upon CAF closure in the flat covered limit (Fig. 3A). Movie S3: Theoretically computed temporal evolution of the CC shape upon CAF closure (Fig. 4A).

## Material and methods

Detailed experimental methods can be found in (17). A short summary is provided in S.I. Theoretical methods are detailed in the S.I.

## Bibliography

1. Andrew D. Rhim, Emily T. Mirek, Nicole M. Aiello, Anirban Maitra, Jennifer M. Bailey, Florencia McAllister, Maximilian Reichert, Gregory L. Beatty, Anil K. Rustgi, Robert H. Vonderheide, Steven D. Leach, and Ben Z. Stanger. EMT and Dissemination Precede Pancreatic Tumor Formation. Cell, 148(1):349–361, January 2012. ISSN 0092-8674, 1097-4172. doi: 10.1016/j.cell.2011.11.025. Publisher: Elsevier.

2. Min Yu, Aditya Bardia, Ben S. Wittner, Shannon L. Stott, Malgorzata E. Smas, David T. Ting, Steven J. Isakoff, Jordan C. Ciciliano, Marissa N. Wells, Ajay M. Shah, Kyle F. Concannon, Maria C. Donaldson, Lecia V. Sequist, Elena Brachtel, Dennis Sgroi, Jose Baselga, Sridhar Ramaswamy, Mehmet Toner, Daniel A. Haber, and Shyamala Maheswaran. Circulating Breast Tumor Cells Exhibit Dynamic Changes in Epithelial and Mesenchymal Composition. Science, 339(6119):580–584, February 2013. ISSN 0036-8075, 1095-9203. doi: 10.1126/science.1228522. Publisher: American Association for the Advancement of Science Section: Report.

3. M. Angela Nieto, Ruby Yun-Ju Huang, Rebecca A. Jackson, and Jean Paul Thiery. EMT: 2016. Cell, 166(1):21–45, June 2016. ISSN 0092-8674, 1097-4172. doi: 10.1016/j.cell.2016.06.028.

4. Leonardo S. Lino-Silva, Rosa A. Salcedo-Hernández, and Armando Gamboa-Domínguez. tumor budding in rectal cancer. A comprehensive review. Contemp Oncol (Pozn), 22(2):61–74, 2018. ISSN 1428-2526, 1897-4309. doi: 10.5114/wo.2018.77043. Publisher: Termedia.

5. Nicola Aceto, Aditya Bardia, David T. Miyamoto, Maria C. Donaldson, Ben S. Wittner, Joel A. Spencer, Min Yu, Adam Pely, Amanda Engstrom, Huili Zhu, Brian W. Brannigan, Ravi Kapur, Shannon L. Stott, Toshi Shioda, Sridhar Ramaswamy, David T. Ting, Charles P. Lin, Mehmet Toner, Daniel A. Haber, and Shyamala Maheswaran. Circulating Tumor Cell Clusters Are Oligoclonal Precursors of Breast Cancer Metastasis. Cell, 158(5):1110–1122, August 2014. ISSN 0092-8674, 1097-4172. doi: 10.1016/j.cell.2014.07.013.

6. A. C. Rogers, D. C. Winter, A. Heeney, D. Gibbons, A. Lugli, G. Puppa, and K. Sheahan. Systematic review and meta-analysis of the impact of tumor budding in colorectal cancer. British Journal of Cancer, 115(7):831–840, September 2016. ISSN 1532-1827. doi: 10.1038/bjc.2016.274. Number: 7 Publisher: Nature Publishing Group.

7. Frances R. Balkwill, Melania Capasso, and Thorsten Hagemann. The tumor microenvironment at a glance. J Cell Sci, 125(23):5591–5596, December 2012. ISSN 0021-9533, 1477-9137. doi: 10.1242/jcs.116392. Publisher: The Company of Biologists Ltd Section: Cell Science at a Glance.

8. Sarah T. Boyle, M. Zahied Johan, and Michael S. Samuel. tumor-directed microenvironment remodelling at a glance. Journal of Cell Science, 133(24), December 2020. ISSN 0021-9533. doi: 10.1242/jcs.247783.

9. Alexandros Glentis, Philipp Oertle, Pascale Mariani, Aleksandra Chikina, Fatima El Marjou, Youmna Attieh, Francois Zaccarini, Marick Lae, Damarys Loew, Florent Dingli, Philemon Sirven, Marie Schoumacher, Basile G. Gurchenkov, Marija Plodinec, and Danijela Matic Vignjevic. Cancer-associated fibroblasts induce metalloprotease-independent cancer cell invasion of the basement membrane. Nature Communications, 8(1):1–13, October 2017. ISSN 2041-1723. doi: 10.1038/s41467-017-00985-8.

10. Youmna Attieh, Andrew G. Clark, Carina Grass, Sophie Richon, Marc Pocard, Pascale Mariani, Nadia Elkhatib, Timo Betz, Basile Gurchenkov, and Danijela Matic Vignjevic. Cancer-associated fibroblasts lead tumor invasion through integrin-beta3–dependent fibronectin assembly. J Cell Biol, 216(11):3509–3520, November 2017. ISSN 0021-9525. doi: 10.1083/jcb.201702033. Publisher: The Rockefeller University Press.

11. Begum Erdogan, Mingfang Ao, Lauren M. White, Anna L. Means, Bryson M. Brewer, Lijie Yang, M. Kay Washington, Chanjuan Shi, Omar E. Franco, Alissa M. Weaver, Simon W. Hayward, Deyu Li, and Donna J. Webb. Cancer-associated fibroblasts promote directional cancer cell migration by aligning fibronectinCAFs direct cancer cell migration via fibronectin. J Cell Biol, 216(11):3799–3816, November 2017. ISSN 0021-9525. doi: 10.1083/jcb.201704053.

12. Cedric Gaggioli, Steven Hooper, Cristina Hidalgo-Carcedo, Robert Grosse, John F. Marshall, Kevin Harrington, and Erik Sahai. Fibroblast-led collective invasion of carcinoma cells with differing roles for RhoGTPases in leading and following cells. Nature Cell Biology, 9 (12):1392–1400, December 2007. ISSN 1476-4679. doi: 10.1038/ncb1658.

13. Anna Labernadie, Takuya Kato, Agustí. Brugués, Xavier Serra-Picamal, Stefanie Derzsi, Esther Arwert, Anne Weston, Victor González-Tarragó, Alberto Elosegui-Artola, Lorenzo Albertazzi, Jordi Alcaraz, Pere Roca-Cusachs, Erik Sahai, and Xavier Trepat. A mechanically active heterotypic E-cadherin/N-cadherin adhesion enables fibroblasts to drive cancer cell invasion. Nature Cell Biology, 19(3):224–237, March 2017. ISSN 1476-4679. doi: 10.1038/ncb3478.

14. Erik Sahai, Igor Astsaturov, Edna Cukierman, David G. DeNardo, Mikala Egeblad, Ronald M. Evans, Douglas Fearon, Florian R. Greten, Sunil R. Hingorani, Tony Hunter, Richard O. Hynes, Rakesh K. Jain, Tobias Janowitz, Claus Jorgensen, Alec C. Kimmelman, Mikhail G. Kolonin, Robert G. Maki, R. Scott Powers, Ellen Puré, Daniel C. Ramirez, Ruth Scherz-Shouval, Mara H. Sherman, Sheila Stewart, Thea D. Tlsty, David A. Tuveson, Fiona M. Watt, Valerie Weaver, Ashani T. Weeraratna, and Zena Werb. A framework for advancing our understanding of cancer-associated fibroblasts. Nature Reviews Cancer, 20 (3):174–186, January 2020. ISSN 1474-1768. doi: 10.1038/s41568-019-0238-1.

15. Berna C. Özdemir, Tsvetelina Pentcheva-Hoang, Julienne L. Carstens, Xiaofeng Zheng, Chia-Chin Wu, Tyler R. Simpson, Hanane Laklai, Hikaru Sugimoto, Christoph Kahlert, Sergey V. Novitskiy, Ana De Jesus-Acosta, Padmanee Sharma, Pedram Heidari, Umar Mahmood, Lynda Chin, Harold L. Moses, Valerie M. Weaver, Anirban Maitra, James P. Allison, Valerie S. LeBleu, and Raghu Kalluri. Depletion of carcinoma-associated fibroblasts and fibrosis induces immunosuppression and accelerates pancreas cancer with reduced survival. Cancer Cell, 25(6):719–734, June 2014. doi: 10.1016/j.ccr.2014.04.005.

16. Daniel Öhlund, Abram Handly-Santana, Giulia Biffi, Ela Elyada, Ana S. Almeida, Mariano Ponz-Sarvise, Vincenzo Corbo, Tobiloba E. Oni, Stephen A. Hearn, Eun Jung Lee, Iok In Christine Chio, Chang-Il Hwang, Hervé Tiriac, Lindsey A. Baker, Dannielle D. Engle, Christine Feig, Anne Kultti, Mikala Egeblad, Douglas T. Fearon, James M. Crawford, Hans Clevers, Youngkyu Park, and David A. Tuveson. Distinct populations of inflammatory fibroblasts and myofibroblasts in pancreatic cancer. Journal of Experimental Medicine, 214(3): 579–596, February 2017. doi: 10.1084/jem.20162024.

17. Jorge Barbazan, Carlos Pérez-González, Manuel Gómez-González, Mathieu Dedenon, Sophie Richon, Ernest Latorre, Marco Serra, Pascale Mariani, Stéphanie Descroix, Pierre Sens, Xavier Trepat, and Danijela Matic Vignjevic. Cancer-associated fibroblasts actively compress cancer cells and modulate mechanotransduction. Nature Communications, 14 (1):6966, November 2023. ISSN 2041-1723. doi: 10.1038/s41467-023-42382-4.

18. Joke Tommelein, Laurine Verset, Tom Boterberg, Pieter Demetter, Marc Bracke, and Olivier De Wever. Cancer-Associated Fibroblasts Connect Metastasis-Promoting Communication in Colorectal Cancer. Front. Oncol., 5, 2015. ISSN 2234-943X. doi: 10.3389/fonc.2015.00063.

19. Olivier Cochet-Escartin, Jonas Ranft, Pascal Silberzan, and Philippe Marcq. Border Forces and Friction Control Epithelial Closure Dynamics. Biophysical Journal, 106(1):65–73, January 2014. ISSN 0006-3495. doi: 10.1016/j.bpj.2013.11.015. Publisher: Elsevier.

20. Sri Ram Krishna Vedula, Grégoire Peyret, Ibrahim Cheddadi, Tianchi Chen, Agustí. Brugués, Hiroaki Hirata, Horacio Lopez-Menendez, Yusuke Toyama, Luís Neves de Almeida, Xavier Trepat, Chwee Teck Lim, and Benoit Ladoux. Mechanics of epithelial closure over non-adherent environments. Nat Commun, 6(1):1–10, January 2015. ISSN 2041-1723. doi: 10.1038/ncomms7111.

21. Simon Begnaud, Tianchi Chen, Delphine Delacour, René-Marc Mège, and Benoît Ladoux. Mechanics of epithelial tissues during gap closure. Current Opinion in Cell Biology, 42: 52–62, October 2016. ISSN 0955-0674. doi: 10.1016/j.ceb.2016.04.006.

22. C. Redon, F. Brochard-Wyart, and F. Rondelez. Dynamics of dewetting. Phys. Rev. Lett., 66(6):715–718, February 1991. doi: 10.1103/PhysRevLett.66.715.

23. T. Vilmin and E. Raphaël. Dewetting of thin polymer films. Eur. Phys. J. E, 21(2):161–174, October 2006. ISSN 1292-895X. doi: 10.1140/epje/i2006-10057-5.

24. Andrew M. J. Edwards, Rodrigo Ledesma-Aguilar, Michael I. Newton, Carl V. Brown, and Glen McHale. Not spreading in reverse: The dewetting of a liquid film into a single drop. Science Advances, 2(9):e1600183, September 2016. ISSN 2375-2548. doi: 10.1126/sciadv.1600183.

25. C. Blanch-Mercader, R. Vincent, E. Bazellières, X. Serra-Picamal, X. Trepat, and J. Casademunt. Effective viscosity and dynamics of spreading epithelia: a solvable model. Soft Matter, 13(6):1235–1243, February 2017. ISSN 1744-6848. doi: 10.1039/C6SM02188C.

26. G. Duclos, C. Blanch-Mercader, V. Yashunsky, G. Salbreux, J.-F. Joanny, J. Prost, and P. Silberzan. Spontaneous shear flow in confined cellular nematics. Nature Phys, 14(7):728–732, July 2018. ISSN 1745-2481. doi: 10.1038/s41567-018-0099-7.

27. Carlos Pérez-González, Ricard Alert, Carles Blanch-Mercader, Manuel Gómez-González, Tomasz Kolodziej, Elsa Bazellieres, Jaume Casademunt, and Xavier Trepat. Active wetting of epithelial tissues. Nature Phys, 15(1):79–88, January 2019. ISSN 1745-2481. doi: 10.1038/s41567-018-0279-5.

28. Carles Blanch-Mercader, Pau Guillamat, Aurélien Roux, and Karsten Kruse. Quantifying Material Properties of Cell Monolayers by Analyzing Integer Topological Defects. Phys. Rev. Lett., 126(2):028101, January 2021. doi: 10.1103/PhysRevLett.126.028101. Publisher: American Physical Society.

29. Nargess Khalilgharibi, Jonathan Fouchard, Nina Asadipour, Ricardo Barrientos, Maria Duda, Alessandra Bonfanti, Amina Yonis, Andrew Harris, Payman Mosaffa, Yasuyuki Fujita, Alexandre Kabla, Yanlan Mao, Buzz Baum, José J. Muñoz, Mark Miodownik, and Guillaume Charras. Stress relaxation in epithelial monolayers is controlled by the actomyosin cortex. Nat. Phys., 15(8):839–847, August 2019. ISSN 1745-2481. doi: 10.1038/s41567-019-0516-6.

30. S. Tlili, J. Yin, J.-F. Rupprecht, M. A. Mendieta-Serrano, G. Weissbart, N. Verma, X. Teng, Y. Toyama, J. Prost, and T. E. Saunders. Shaping the zebrafish myotome by intertissue friction and active stress. PNAS, 116(51):25430–25439, December 2019. ISSN 0027-8424, 1091-6490. doi: 10.1073/pnas.1900819116. Publisher: National Academy of Sciences Section: PNAS Plus.

31. Charlotte Alibert, Bruno Goud, and Jean-Baptiste Manneville. Are cancer cells really softer than normal cells? Biology of the Cell, 109(5):167–189, 2017. ISSN 1768-322X. doi: 10.1111/boc.201600078. _eprint: https://onlinelibrary.wiley.com/doi/pdf/10.1111/boc.201600078.

32. Zheng Peng, Meng Hao, Haibo Tong, Hongmei Yang, Bin Huang, Zhigang Zhang, and Kathy Qian Luo. The interactions between integrin α5β1 of liver cancer cells and fibronectin of fibroblasts promote tumor growth and angiogenesis. International Journal of Biological Sciences, 18(13):5019–5037, 2022. ISSN 1449-2288. doi: 10.7150/ijbs.72367.

33. Gonca Erdemci-Tandogan and M. Lisa Manning. Effect of cellular rearrangement time delays on the rheology of vertex models for confluent tissues. PLOS Computational Biology, 17(6):e1009049, June 2021. ISSN 1553-7358. doi: 10.1371/journal.pcbi.1009049.

